# Nutraceutical profiles of apricots (*Prunus armeniaca* L.) as a source of fruit quality traits for breeding

**DOI:** 10.1101/2020.11.30.403527

**Authors:** Helena Gómez-Martínez, Almudena Bermejo, María Luisa Badenes, Elena Zuriaga

## Abstract

In a social context of increasingly concern about healthy diets, the development of new varieties with enhanced content in nutraceutical compounds is an increasingly important objective of the fruit breeding programs currently developed. In this sense, apricot is a fruit crop very appreciated by consumers due to its organoleptic characteristics, but also plays an important role in human nutrition due to its contain of phytocompounds as sugars, organic acids, vitamins and polyphenols. In this work, new selections from the apricot breeding program carried out at the Instituto Valenciano de Investigaciones Agrarias (IVIA) and traditional varieties have been analysed aimed at identifying sources of genetic variation for fruit quality. For this purpose, sugar content, organic acids and ascorbic acid were studied during two crop years. Results revealed sucrose and glucose as the major sugars, malic and citric acid as the main organic acids, and diverse ascorbic acid content among the cultivars studied. Results obtained pointed some accessions as potential sources to increase fruit quality. In addition, the study showed that apricot peel is an excellent source of nutraceutical compounds. Moreover, this study opens up new possibilities for future work to study the genetic control of these traits in apricot.

## 1. Introduction

The increasing demand for safe, healthy and nutritious food by consumers, turn the internal quality of the fruit into one of the main goals of the food industry. In this sense, plants and some fruits become a useful source of compounds with a relevant role in improving health (Vieira da Silva *et al*., 2016; Slavin and Lloyd, 2012). In fact, plant extracts and their bioactive compounds are used by the industry to produce functional food (Azmir *et al*., 2013). For this reason, those fruits with high content of these compounds are of high interest for the industry. In this sense, nutraceutical profiles can be used for promotion of fruit consumption as a natural functional food.

Apricot (*Prunus armeniaca* L.) is a stone fruit crop species with a large tradition in the Mediterranean basin countries. World apricot production reached 3.84 million tonnes in 2018, being Turkey, Uzbekistan, Iran, Algeria, Italy and Spain as main producers (http://www.fao.org/faostat/). Despite its wide geographical spread apricot has very specific ecological requirements, so each region usually grows locally adapted cultivars. Significant breeding efforts have been undertaken (Zhebentyayeva *et al*., 2012), leading to a rich diversity apricot germplasm in terms of fruit morphology, harvest season or biotic and abiotic resistances. Apricots are consumed in multiple and diverse ways, including fresh or processed fruits (as dried, canned, jam, juice or even liquors), and also used the apricot kernel oil for medicinal purposes (Zhebentyayeva *et al*., 2012). Apricots are an important source of sugars, fiber, proteins, minerals and vitamins (Sochor *et al*., 2010; Moustafa and Cross, 2019). However, pomological and nutraceutical properties depend on varieties, cultivation systems, fruit storage conditions or developmental stages (Ruiz *et al.*, 2005).

In terms of fruit consumption, organoleptic characteristics are one of the main factors for consumers decision. Notwithstanding, nutraceutical compounds interact with each other and influence the quality properties making it difficult to handle. For instance, the flavour is provided by sucrose, malic acid and volatiles (Xi *et al*., 2016), being sugar and organic acid balance relevant for sweetness. From them, fructose and sucrose are the prominent contributors to sweetness, being the most important sensory quality for consumer satisfaction (Fan *et al*.,2017). Similar results have been found in peach, whose sweetness depends on the overall sugar amount as well as in the specific relative amount of each individual sugar (Kroger *et al*., 2006). Regarding the apricot nutraceutical profile, previous studies have also found glucose and sucrose as the major sugars in both pulp and peel (Xi *et al*., 2016). Moreover, during the fruit ripening a high number of molecular and metabolic changes occur that have a relevant effect in fruit properties (D’Ambrosio *et al*., 2013; Karlova *et al*., 2014; Osorio *et al*., 2013; Seymour *et al*., 2013). For instance, organic acids increase during the early stages of fruit development and decrease when fruits were full-ripped, being malic the most important organic acid in apricot (Xi *et al*., 2016). Additionally, fruits and vegetables constitute the main source of ascorbate in the human diet, so rising its content in highly consumed fruits would clearly have an impact on human nutrition (Fenech *et al*., 2019). Moreover, ascorbate content has been also related with elevated stress tolerance (Fenech *et al*., 2019). In fact, foliar application of ascorbic acid on peach trees resulted in improving the yield and fruit quality (Sajid *et al*., 2017). Previous studies found that vitamin C content in apricot could reach up to 100 mg/100 g dry weight (Akin *et al*., 2008), showing the potential of this species as a source of this vitamin.

In conclusion, apricot germplasm represent a diverse source of phytocompounds that can be exploited for breeding purposes in order to develop new varieties with higher content of these nutraceutical compounds. The apricot breeding program at the Instituto Valenciano de Investigaciones Agrarias (IVIA) has the purpose of obtaining new varieties, with high fruit quality, resistant to the Plum Pox virus (PPV), self-compatibles and well-adapted to the Southern European environment (Martínez-Calvo et al., 2009). PPV is the main limiting factor for apricot production worldwide, hence, during the last decades, development of PPV resistant varieties has been the main objective of almost any apricot breeding program. However, for this purpose, just some North American cultivars not well-adapted to Mediterranean conditions were identified and used as resistance donors (Martínez-Gómez *et al*., 2000). This represents a challenge especially in the current climate change scenario affecting the Mediterranean basin, with increasingly mild winters.

The objective of the present work is to assess the fruit quality characterization of 1 North-American, 3 Spanish (Valencian Community) and 9 accessions from the IVIA’s apricot breeding program aimed at identifying the most convenient genotypes for increasing the fruit quality of apricot while keeping the adaptability to warm winters. In this study we analyse sugars (sucrose, fructose, and glucose), ascorbic acid, and organic acids (citric, malic, succinic and fumaric).

## 2. Material and Methods

### 2.1. Plant Material

Thirteen apricot genotypes were used, including 3 well-known cultivars from the Mediterranean Basin, 1 North-American, and 9 selections from the IVIA’s breeding program resistant to PPV (Table 1). All of them are kept at the collection of the IVIA in Moncada (Valencia, Spain). Five fruits per tree were harvested at the ripening stage during 2 growing seasons (2016-2017). For each fruit, the peel was separated from the flesh with a peeler. A mix of 5 fruits (peel or flesh, respectively) was frozen with liquid nitrogen and kept at −80°C until processing. Peel samples were freeze-dried and powdered. Tissue homogenization was carried out using a Polytrom 3100 (Kinematica AG, Switzerland) and a vortex for the flesh and peel samples, respectively.

**Table 1:** Apricot accessions analysed

| **Genotype** | **Pedigree** | **Origin** | **Harvest date** |  |
| --- | --- | --- | --- | --- |
|  |  | **Country** | **2016** | **2017** |
| Goldrich | Sunglo x Perfection | USA | June 22nd | 9th June |
| Canino | Unknown | Spain (Valencia) | June 3rd | 31st May |
| Mitger | Unknown | Spain (Valencia) | June 3rd | 25th May |
| Tadeo | Unknown | Spain (Valencia) | June 15th | 9th June |
| Dama Rosa | Goldrich x Ginesta | Spain (IVIA) | June 6th | 9th June |
| Dama Taronja | Goldrich x Katy | Spain (IVIA) | June 10th | 9th June |
| GG9310 | Goldrich x Ginesta | Spain (IVIA) | June 6th | 9th June |
| GG979 | Goldrich x Ginesta | Spain (IVIA) | June 13th | 9th June |
| GP9817 | Goldrich x Palau | Spain (IVIA) | June 13th | 9th June |
| HG9821 | Harcot x Ginesta | Spain (IVIA) | June 8th | 25th June |
| HG9850 | Harcot x Ginesta | Spain (IVIA) | June 3rd | 25th June |
| HM964 | Harcot x Mitger | Spain (IVIA) | June 1st | 2nd June |
| SEOP934 | SEO x Palau | Spain (IVIA) | June 8th | 2nd June |

### 2.2. Sample processing and HPLC analysis

For sample processing, 1 g of flesh or 10-20 mg of freeze-dried peels were mixed with 1.5 mL of 5% metaphosphoric acid solution, 1 mL of water of LC-MS grade and 1 mL of 0.1% H_2_SO_4_ solution for ascorbic acid, sugars and organic acids extraction, respectively. Then the sample was homogenized and centrifuged at 4°C for 20 min at 8.050xg.

Compounds were identified on the bases of comparing their retention times, UV-vis spectra and mass spectrum data with authentic standards obtained from Sigma-Aldrich using an external calibration curve. In addition, standards were run daily with samples for validation. All the solvents used were of LC-MS grade. Three samples per cultivar were analysed and all the samples were run in triplicate. The Empower 2 software (Waters, Spain) was used for data processing.

#### 2.2.1. Ascorbic acid

Total ascorbic acid was extracted according to the method previously described by Cano *et al*. (2011) adapted to a microliter format (Sdiri *et al.*, 2012) and using *DL*-dithiothreitol (DTT) as reducing reagent of dehydroascorbic acid to ascorbic acid. After centrifugation, 1mL of supernatant was mixed with 200 μL of DTT (20 mg/mL) and maintained for 2 h in the dark, then filtered through 0.45 μm filter. It was analysed by HPLC-DAD in an Alliance liquid chromatographic system (Waters, Barcelona, Spain) equipped with a 2695 separation module coupled to a 2996 photodiode array detector, and a reverse-phase C_18_ column Tracer Excel 5 μm 120 OSDB (250 mm × 4.6 mm) (Teknokroma, Barcelona, Spain) with an isocratic mobile phase of methanol:0.6% acetic acid (5:95) at a flow rate of 1 mL/min, and injection volume was 5 μL. The quantification was performed at 245 nm.

#### 2.2.2. Sugars

Sucrose, glucose, and fructose were extracted as described by Sdiri *et al.* (2012). After centrifugation, samples were filtered through a 0.45 μm nylon filter and analysed by an HPLC system equipped with a Waters 515 HPLC pump, a Waters 2414 refractive index detector, a 5-μm Tracer Carbohydr column (250 mm × 4.5 mm) (Teknokroma, Barcelona, Spain), and a 20-μL loop Rheodyne injector were used for the sugar analysis. The mobile phase was composed of acetonitrile and water (75:25) at a flow rate of 1.0 mL/min.

#### 2.2.3. Organic acids

Citric, malic, succinic and fumaric acids were extracted as described by Sdiri *et al*. (2012). After centrifugation, the supernatant was filtered through a 0.45 μm filter, analysed by HPLC-DAD and confirmed by HPLC-MS under electrospray ion negative conditions using a ZQ2000 mass detector. The sample temperature was 5°C and column temperature was 35°C. Capillary voltage was 3.0 kV, cone voltage was 23 V, source temperature was 100°C, desolvation temperature was 200°C and desolvation gas flow was 400 L/h. Full data acquisition was performed by scanning from 100 to 400 uma in the centroid mode. An ICSep ICE-COREGEL 87H3 column (Transgenomic, UK), an ICSep ICE-COREGEL 87H guard kit, and an automatic injector were used for chromatographic separation. The solvent system was an isocratic mobile phase of 0.1% H_2_SO_4_ solution. The total run time was 20 min at 0.6 mL/min, and the injection volume was 5 μL.

### 2.3 Data analysis

Data were analysed with R (R Core Team, 2012) using R-studio software (v.3.5.3) with *stats*, *ggbiplot, readxl, graphics* and *grDevices* packages. Normality and homoscedasticity were checked using Shapiro-Wilk and Bartlett tests respectively. Next, the non parametric Kruskal-Wallis test was used to make all samples comparisons. Correlation coefficients among the variables were determined using the Spearman method. Principal component analysis (PCA), using centered and scaled data, was conducted to visualize the relationships between accessions and variables.

#### 2.3.1. Sweetness Index (SI) and Total Sweetness Index (TSI)

In order to determine the sweetness perception of fruits, both index were calculated according to Magwaza and Opara (2015) following the equations:

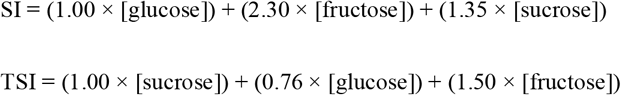

## 3. Results

### 3.1 Sugars

Fructose, glucose and sucrose content in peel and flesh showed significant differences (p≤ 0.05) between the accessions analysed (Figure 1, Table S1). Regarding total sugar content in flesh, SEOP934 showed the highest value (12.34 g/100 g FW) and HG9850 the lower one (5.75 g/100 g FW) (Figure 1A). In all cases sucrose was the predominant sugar, ranging from 65.1 to 90.3% of the total. For each sugar, Tadeo showed the highest content of fructose (0.48 g/100 g FW) and glucose (3.11 g/100 g FW), and SEOP934 showed the highest quantity of sucrose (10.3 g/100 g FW). Regarding peel content, Kruskal-Wallist test showed an effect of the crop year over the fructose and glucose content, but not over sucrose. According to the Spearman correlation analysis, seven significant correlations were observed between the analysed sugars (Figure 2, Table S2). Mainly, fructose and glucose appear positively correlated between tissues and also between years, while in peel fructose peel appeared negatively correlated with sucrose. The North American Goldrich cultivar showed the lower total sugar content in 2016 (38.19 g/100 g DW) and the second lowest in 2017 (23.68 g/100 g DW), mainly due to its low sucrose content (12.41 and 4.63 g/ 100 g DW, respectively). In general, the well-known cultivars from the Mediterranean Basin showed high sugar content and the accessions belonging to the IVIA’s breeding program showed an intermediate content between them and Goldrich. For each sugar, fructose ranged between the 5.9-21.2 % of total sugar measured, glucose between 19.7-45.6% and sucrose between 35.5-71.1%. As a measure of sweetness, SI and TSI index were calculated (Table 2). According to these indexes, fruits with identical total sugar content but with relatively more fructose or sucrose will taste sweeter. Overall, the Spanish cultivar Tadeo and 2 of the selections of the breeding program, HG9821 and SEOP934, were the sweetest according to these indexes. Contrary, the selections Dama Rosa, GG979 and HG9850 have the lower values.

**Table 2:**
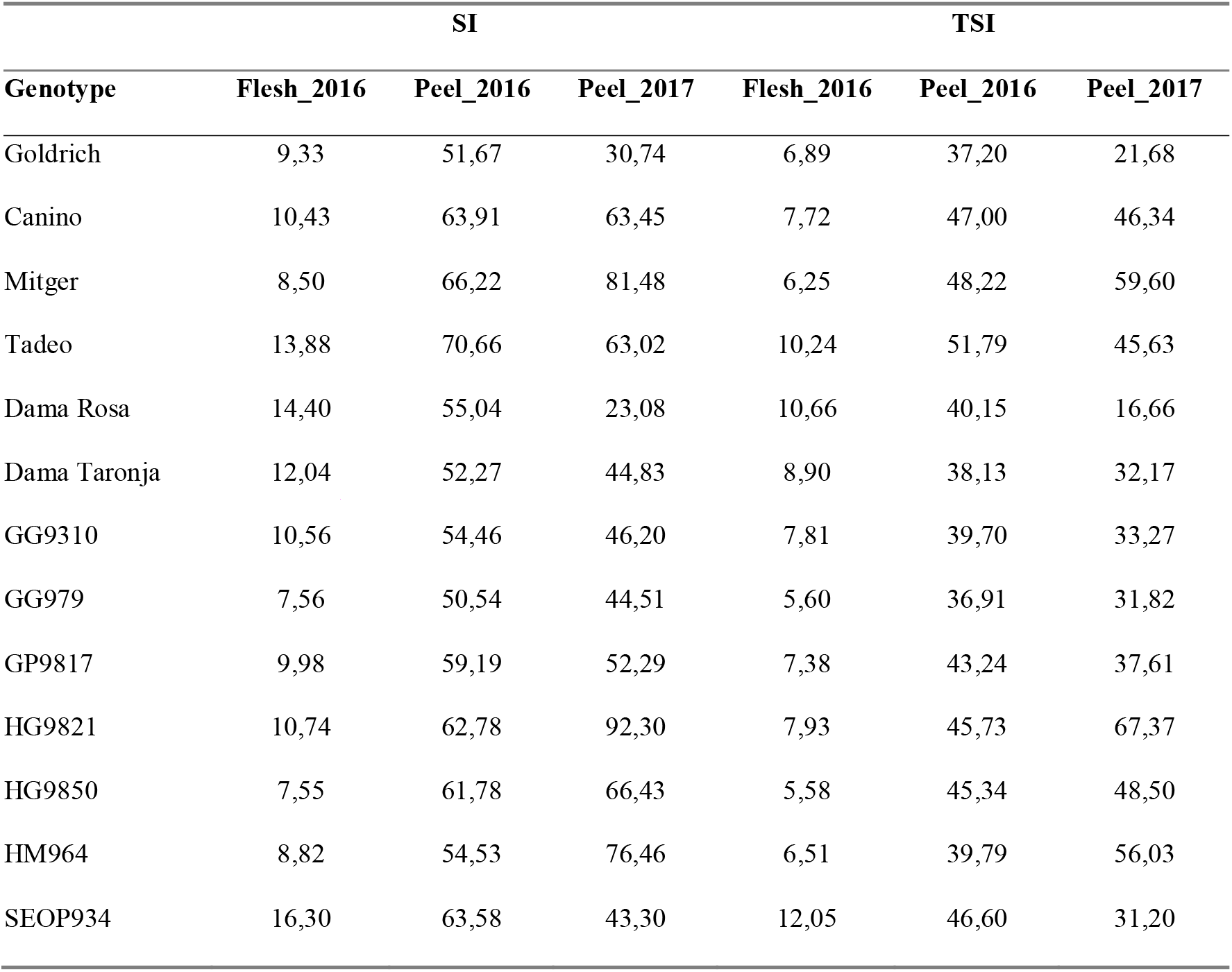
Sweetness estimation, SI and TSI were calculated according to Magwaza and Opara (2015).

**Figure 1:**
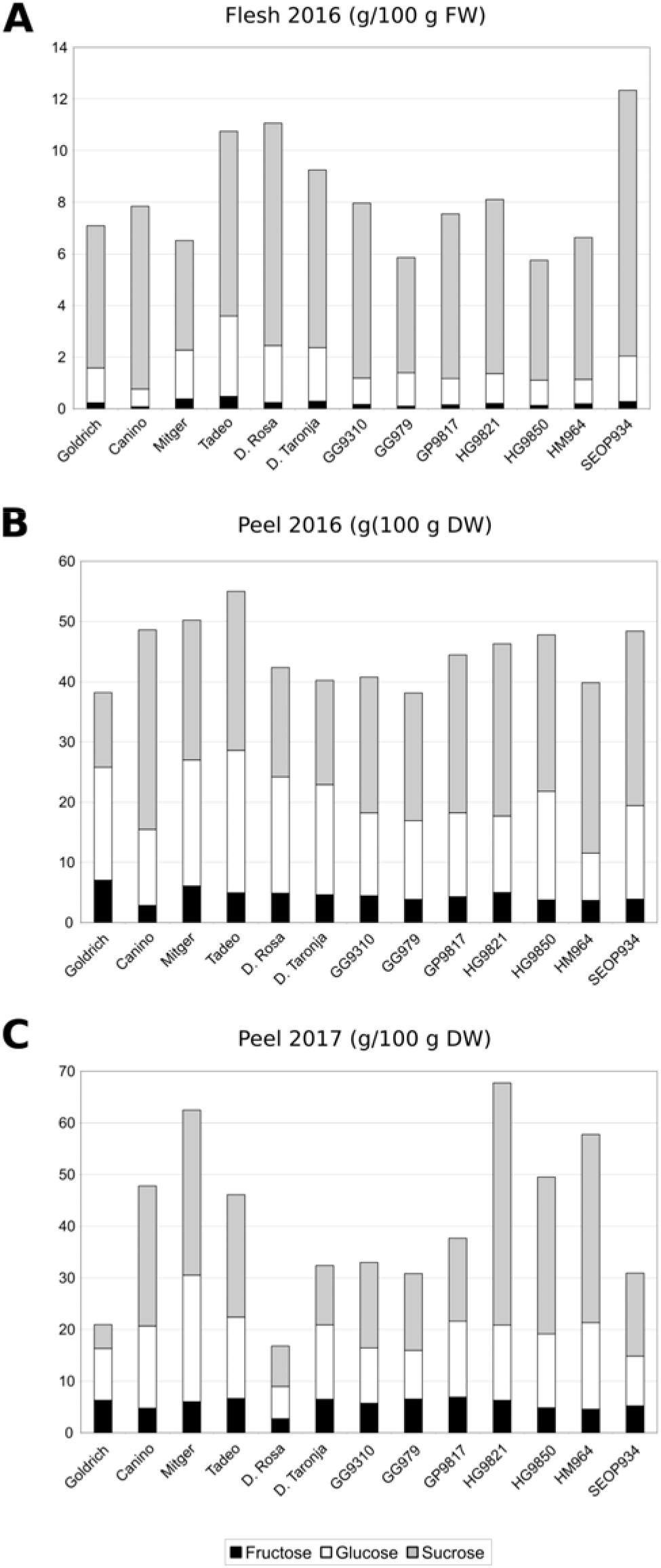
Profiles of sugar content in flesh (g/100 g fresh weight (FW)) and peel (g /100 g dry weight (DW)) during 2016 and 2017.

**Figure 2:**
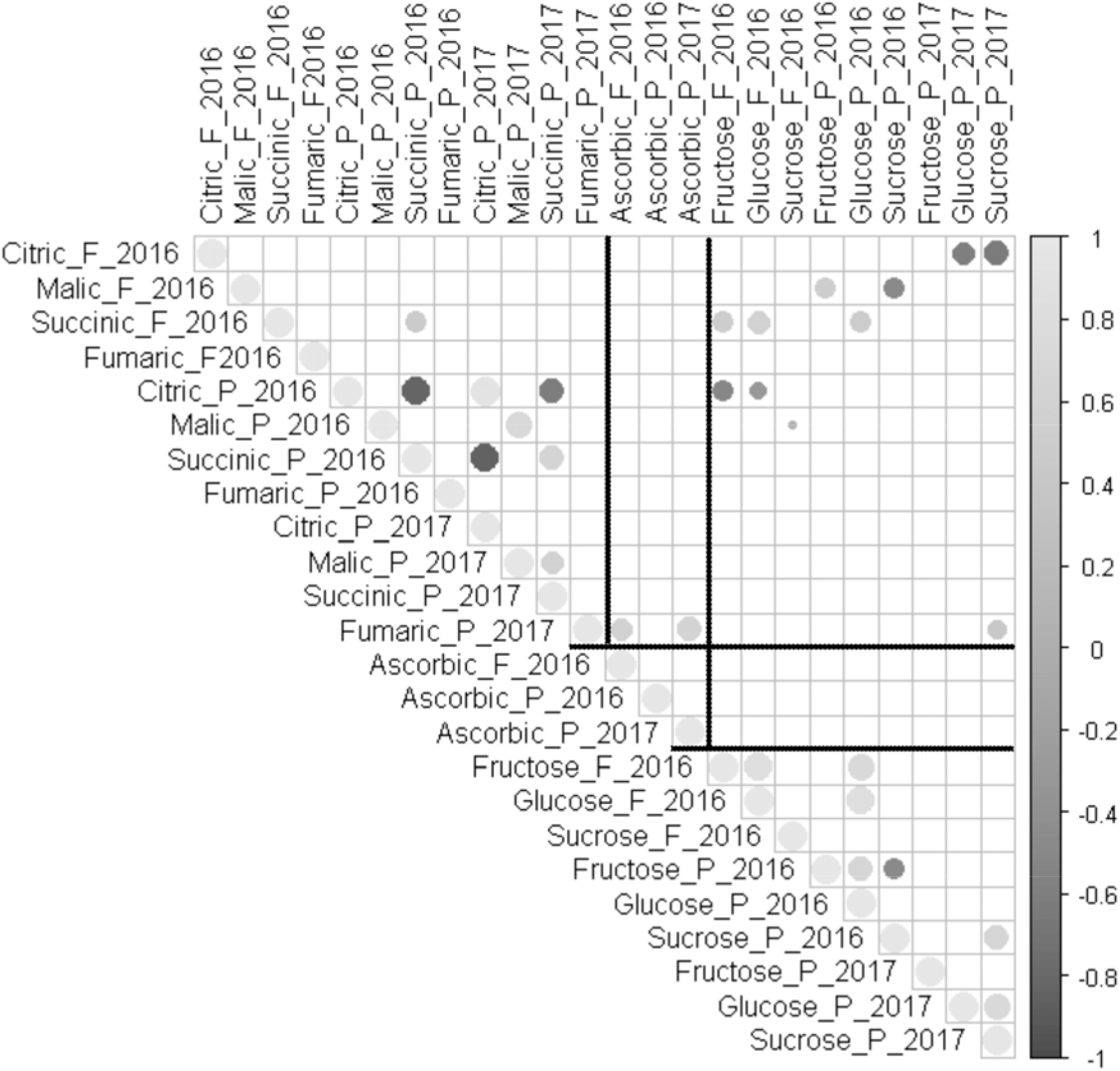
Significant correlations between variables analysed (α =0,05)

### 3.2. Organic acids

Significant differences among the apricot accessions were observed for citric, malic, succinic and fumaric acids content (Figure 3, Table S3). Citric acid was the main organic acid in flesh in all cases, ranging from 50-80.1%. Malic acid was the second one except for Tadeo and SEOP934. Succinic represents between 7-22.93% of the organic acids measured and fumaric just between 0.07 and 0.17%. Regarding total content in flesh, Dama Rosa showed the highest value (3.254 g/100 g FW) and Mitger the lower one (1.562 g/100 g FW) (Figure 3A). In this case, 8 significant correlations were detected between organics (Figure 2), being the most notorious the negative correlation between succinic and citric acids in peel. As in the case of sugars, an effect of crop year was observed in the fumaric acid peel content, with some accessions showing higher values in 2017 (Figure 3B and 3C). Citric acid was the main organic acid in peel in all cases except for Mitger (~14.5%%) and Tadeo (~20%), which showed both years a higher content of malic acid (around 62% and 55%, respectively), and Dama Rosa with 46.6% in 2017. Succinic acid was the third one, except for Tadeo (~22.2%), which showed more content than malic (~20%). Regarding the total content in peel, Goldrich showed the highest values (20.7 and 31.7 g /100 g DW), and Tadeo the lower ones (9.6 and 11.3 g/100 gDW).

**Figure 3:**
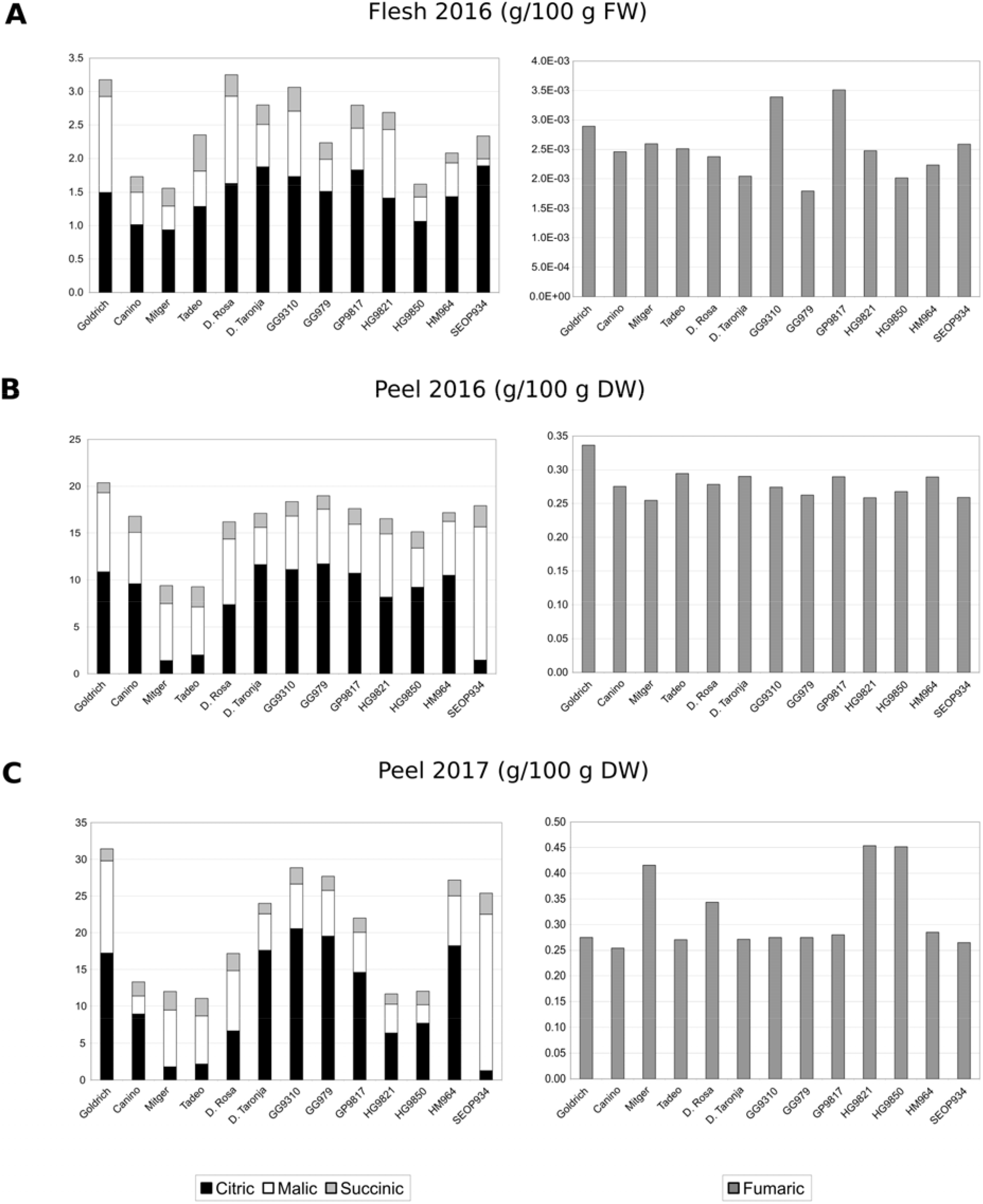
Profiles of organic acids content in flesh (g/100 g FW) and peel (g /100 g DW) during 2016 and 2017.

### 3.3 Ascorbic acid

Results of ascorbic acid content in peel and flesh of the genotypes studied in the two crop years are in Figure 4 and Table S4. Significant differences were found among genotypes and crop years (p-value = 0,0026). In flesh, values ranged from 9.11 mg/100 g FW (SEOP934) and 13.08 mg/100 g FW (HG9821). Regarding peel content, Mitger and HG9850 showed the highest values in 2016 (185.02 mg/100 g DW) and 2017 (192.82 mg/100 g DW) respectively. In general, the accessions belonging to the IVIA’s breeding program showed equal or higher content than Canino, which showed the lowest value from the Mediterranean cultivars.

**Figure 4:**
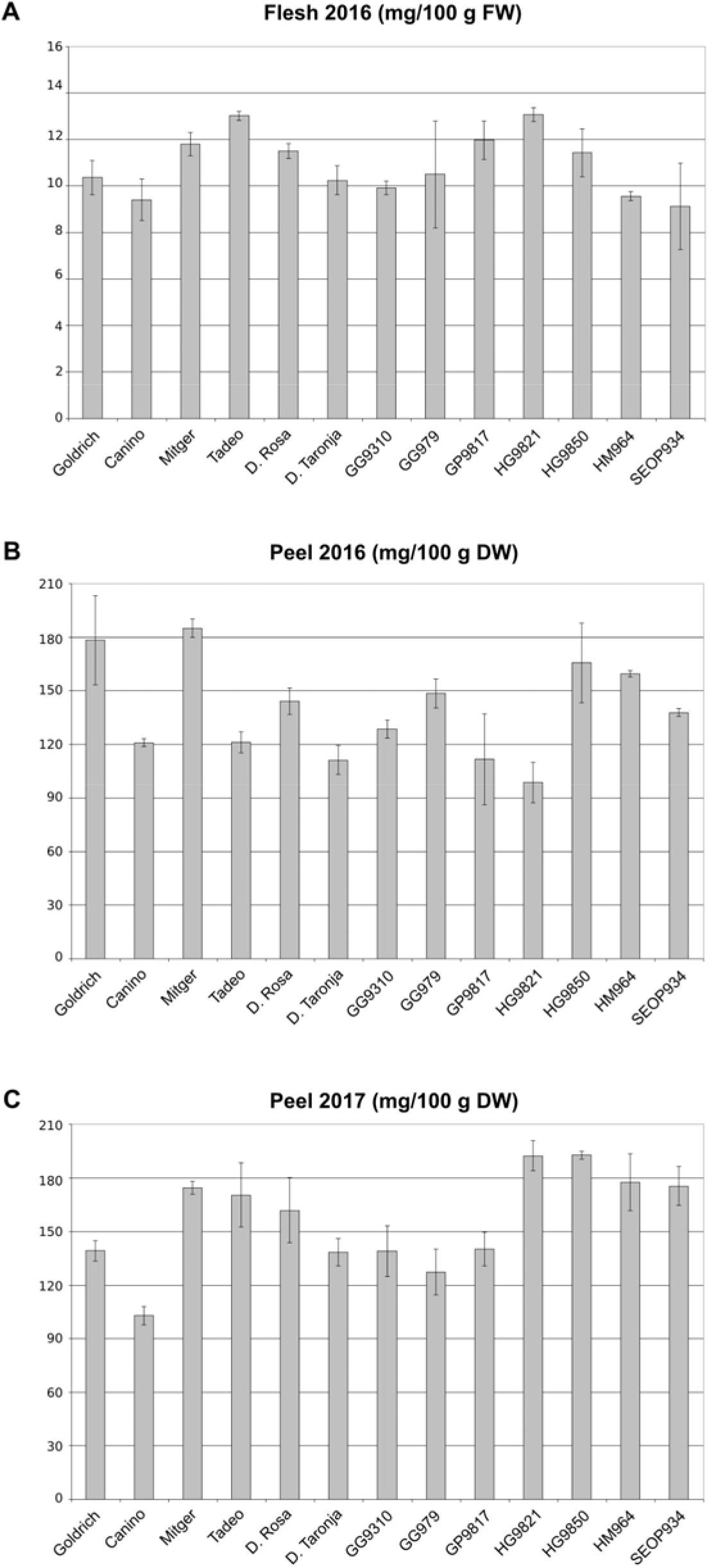
Ascorbic acid content in flesh (mg/100 g FW) and peel (mg /100 g DW) during 2016 and 2017.

### 3.4. Correlations and Principal Component Analysis

As consumer preferences are highly influenced by the balance of sugar and organic acids content, relations between all the analysed compounds were also studied (Figure 2). For instance, citric content in flesh was negatively correlated with glucose and sucrose peel content, while in peel was negatively correlated with fructose and glucose flesh content. Malic content in flesh was positively correlated with fructose content and negatively with sucrose, both in peel. Succinic content in flesh was positively correlated with fructose and glucose. Finally, fumaric acid showed positive correlation with ascorbic acid content.

In order to explore the variability observed in the accessions, the nutraceutical compounds analysed each year were submitted to principal component analyses (PCA). As results with each independent data sets were quite similar, just the PCA for 2017 is shown (Figure 5). First 3 principal components (PC1, PC2 and PC3) accounted for 78.1% of the total variance (36.5%, 27.4% and 14.2%, respectively). PC1 was positively correlated mainly with sucrose, fumaric acid and ascorbic, while negatively with citric acid content. PC2 showed a positive correlation with citric acid and fructose, and negatively with malic and succinic acids. The distribution of the accessions in the space of the 3 first components showed one group of 6 accessions (Goldrich, GG979, GP9817, GG9310, Dama Taronja and Canino) situated at negative values of PC1 and positive values of PC2. Rest of the accessions appear more distributed, and just two of them were close to each other, HG9821 and HG9850, both descendants from the same cross.

**Figure 5:**
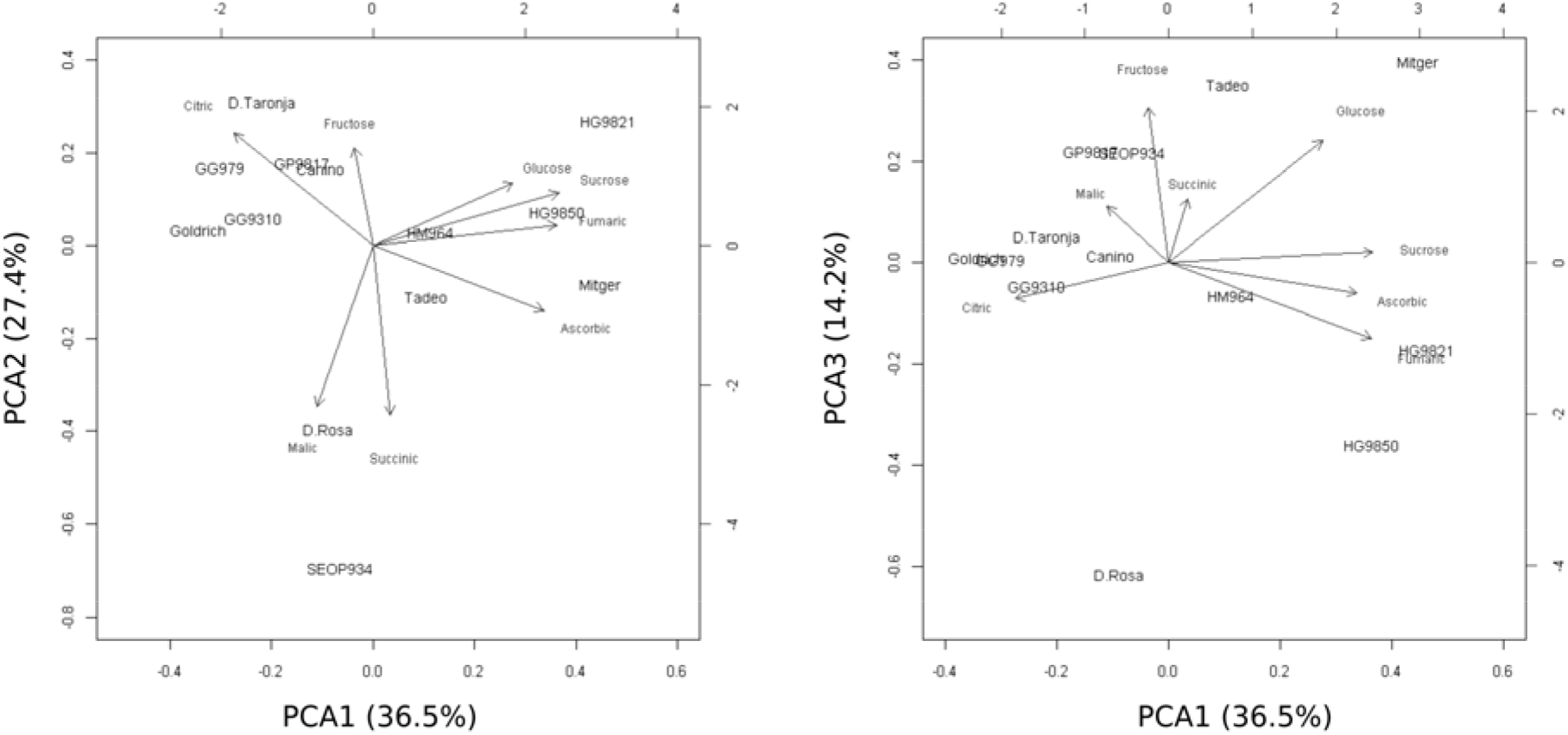
Principal Component Analysis for 2017 data. A: Representation of the accessions in the space defined by the first and second components; B: Representation of the accessions in the space defined by the first and third components.

## 4. Discussion

Traditionally, plant breeding goals have been focused on yield, stress resistance and external quality traits as appearance and shelf-life. However, consumers are increasingly demanding high quality food. As an example, huge efforts are in progress to recover the lost flavour in tomato cultivars (Tieman *et al*., 2017). Nowadays, internal quality traits have been incorporated as objectives of almost any plant breeding program. The IVIA’s apricot breeding program started in 1993 and was initially focused on introgression of sharka resistance into locally grown cultivars (Martínez-Calvo *et al*., 2009). However, just a handful of North American apricot PPV resistant cultivars, adapted to cold-growing conditions, have been identified (Martínez-Gómez *et al*., 2000). Despite the crosses with those cultivars introduce also undesirable traits, the hybrids obtained in the breeding program represent a good opportunity to incorporate new breeding goals and to accelerate the development of new varieties better adapted to the Mediterranean basin conditions. In this sense, the characterization of the nutraceutical properties of these germplasm collection allows to identify putative promising accessions and to optimize the design of the future crosses. This study opens also future work to study the genetic control of these traits in apricot. In this work, we analysed 13 accessions of the IVIA’s collection in order to identify the main source of variation for each phytocompound of interest: sugars (sucrose, fructose and glucose), organic acids (citric, malic, succinic and fumaric) and vitamin C (ascorbic acid).

Apricot fruits are a good source of sugars, fiber, proteins, minerals and vitamins (Moustafa and Cross, 2019). Fruit taste is highly dependent of the soluble solids content, which is the sum of sugars, acids and other minor components, however sugars represent the most important proportion. As described in apricot and other *Prunus* species, sucrose, glucose and fructose are the main sugars present in fruits (Bassi and Selli, 1990; Cirilli *et al*., 2016). For instance, sucrose is the predominant sugar (40-85%) in peach, followed by fructose and glucose in variable ratios (Cirilli *et al*., 2016), similarly to our data presented here. According to Bae *et al*. (2014), the content of glucose and fructose was higher than sucrose and sorbitol during fruit growth, these authors also pointed that sucrose increase as major sugar in apricot and plum at the end of maturity, that is in accordance with our results. Consumer perception of sweetness intensity depends on the overall sugar amount but also the specific profile (Cirilli *et al*., 2016). For this sweetness estimation, the contribution of each carbohydrate is calculated, based on the fact that fructose and sucrose are sweeter than glucose (Magwaza and Opara, 2015). Although comparisons with other previous works are complicated for this type of traits, our values are similar to the ones obtained by Fan *et al*., (2017) analyzing northwest Chinese apricots. According to our study, SEOP934, HG9821 and HG9850 could be good candidates as swetness source.

Organic acids also have an important role, with sugars, on apricot taste (Xi *et al*., 2016). All organic acids increase at first and then fall throughout fruit development and ripening process (Xi *et al*., 2016). In agreement with the previous studies already cited, malic and citric acids were predominant in the apricot genotypes analysed. In terms of taste Dolenc-Sturm *et al*., (1999) pointed the stronger acidic taste of malic compared with citric acid, and conclude that the optimal ratio between malic and citric acid is near the value of 0.8. Interestingly, some accessions showed the malic:citric ratio around this value, like Dama Rosa and HG9821, two accessions from the IVIA’s breeding program, and also Goldrich. Interestingly, the PPV resistant Dama Rosa cultivar has been already registered (Badenes *et al*., 2018). Moreover, cultivars with high content in acids and low in sugars could be more appreciated, particularly those with higher citric acid concentration (Dolenc-Sturm *et al*., 1999). Moreover, cultivars with high content of organic acids could be also used as source of these compounds, as they can be used to provide acidity and sour flavour as additive in food products. For instance, malic acid is used for elaboration of sweets and fumaric acid is used as acidulant and antioxidant in soft drinks and cake mixes (Moldes *et al*. 2017). In this sense, several of the selections studied could be useful for the food-industry, like GG9310, GG979, and SEOP934 that appear as good candidates as they showed high contents of total organics acids.

Finally, the ascorbic acid is one of the most important vitamin in fruits (Lee and Kader, 2000), because of its protective activity as antioxidant (Rice-Evans *et al*., 1997). We found significant differences in ascorbic acid contents between crop years and among genotypes. Our results are in agreement with others studies on apricot varieties (Akin *et al*., 2008; Gündogdu *et al*., 2013), with values ranging from 98.70 to 192.82 mg/100g DW among varieties and crop year. HG9850 and HM964 could be suggested as promising cultivars for ascorbic acid content improving due to their high content and stable behaviour in both years.

The increasing demand of healthy products has raised the need of using alternative supplements and additives in food and, fruit nutraceutical compounds can be a good choice since they can be extracted from natural sources and can provide extra health benefits (Moldes *et al*., 2017). Our results suggest that apricot peel is a good source of sugars, vitamins and organic acids, being an interesting provider of nutraceutical compounds. Our results are in agreement with other authors that pointed the apricot peel as an extraordinary source of nutraceutical compounds and an optimum tissue for studying mechanisms of flavour quality formation in fruit (Xi *et al*., 2016; Voo *et al*., 2012). Similar results were found in previous apricot studies (Ruiz *et al*., 2005) and other fruits species like pear (Li *et al*., 2014) or peach (Campbell *et al*., 2013).

## 4. Conclusions

A set of selections and genitors from the IVIA’s apricot breeding collection has been characterized from a nutraceutical point of view and the main sources of variation of the group of genotypes have been identified, which can be considered as a previous step for further breeding. Our results confirmed the diversity among the set of apricot studied regarding to sugars, organic acids and ascorbic acid content.

## Acknowledgments

This study was funded by the Instituto Nacional de Investigación y Tecnología Agraria y Alimentaria (INIA)-FEDER (RTA2017-00011-C03-01) and the Generalitat Valenciana (GV/2016/189). HGM was funded by a fellowship from INIA.

## Declaration of Competing Interest

The authors declare no conflict of interest. The funders had no role in the design of the study; in the collection, analyses, or interpretation of data; in the writing of the manuscript, or in the decision to publish the results.

## Author contributions

Helena Gómez-Martínez: Data curation, Formal analysis, Writing - review & editing; Almudena Bermejo: Data curation, Methodology; María Luisa Badenes: Conceptualization, Funding acquisition, Supervision, Writing - review & editing; Elena Zuriaga: Conceptualization, Data curation, Formal analysis, Funding acquisition, Writing - original draft, Writing - review & editing

---

Table S1: Profiles of sugar content in flesh (g/100 g fresh weight (FW)) and peel (g /100 g dry weight (DW)) during 2016 and 2017.

Table S2: Correlations between variables analysed.

Table S3: Profiles of organic acids content in flesh (g/100 g FW) and peel (g /100 g DW) during 2016 and 2017.

Table S4: Ascorbic acid content in flesh (mg/100 g FW) and peel (g /100 g DW) during 2016 and 2017.

## Postal address

Citriculture and Crop Production Center, Instituto Valenciano de Investigaciones Agrarias (IVIA), CV-315, Km. 10.7, Moncada, 46113 Valencia, Spain

## Notes

### Competing Interest Statement

The authors have declared no competing interest.

